# Programmed Manipulation of RNA Targets By Human Argonaute 2

**DOI:** 10.64898/2026.02.01.703109

**Authors:** Robert F. Lusi, Salvador Moncayo von Hase, Corey Model, Steven M. Banik

## Abstract

Nucleic acid manipulation using programmable ribonucleoprotein complexes (RNPs) has enabled transformative research tools and led to new therapeutic strategies. RNA directly regulates diverse cellular processes,^1^ is a crucial mediator of protein synthesis, and offers advantages in therapeutic targeting and fundamental discovery complementary to those of DNA.^2^ ISC-^3^ and Cas-based^4,5^ scaffolds where the RNP is fused to an effector protein can alter RNA sequence, structure, and function. However, the non-human origins underlying these systems create challenges in therapeutic translation and the presence of non-native proteins can have unintended and little understood effects on cells.^6–8^ Systematic repurposing of human proteins, which have been optimized in the cellular environment by evolution, for expanded programmable functions could reveal new biological principles and bypass the limitations of foreign proteins. Here, we demonstrate that the catalytic engine of the RNA-interference (RNAi) pathway, human Argonaute 2 (AGO2),^9,10^ can be repurposed as a modular targeting domain, and when fused to a C-to-U deaminase, enable AGO-Led Targeted Editing of RNA (ALTER). Using guide RNAs which remodel target RNA structure for selective editing and reduced nuclease activity, we show that ALTER can act on a variety of target transcripts including endogenous mRNAs and lncRNAs, with activities comparable or exceeding those of Cas-based systems.^11,12^ Despite its human origin and role in RNAi, transcriptome-wide RNAseq revealed lower levels of off-target editing compared to Cas-based editing systems. These results demonstrate that AGO2 can be rationally redirected from RNAi to a broader spectrum of RNA manipulations, establishing that intact human proteins can be reconfigured for expanded molecular function.

## Introduction

Programmable small RNA-protein complexes offer the flexibility to convert a single protein into a scaffold for diverse manipulations. Small RNAs used as guides benefit from increased stability from binding to their protein partner and offer unparalleled ease of target programming and chemical modification.^13^ The sequence-specific manipulation of RNA by RNA-guided effectors has emerged as a ubiquitous mechanism for post-transcriptional regulation in prokaryotes and eukaryotes.^9,14^ Cas13/Type VI CRISPR systems derived from prokaryotes and more broadly, OMEGA-family proteins, have been imported into mammalian cells to generate programmable RNA-modulating proteins. Catalytically active Cas13s perform targeted RNA cleavage,^15^ while nuclease-deactivated dCas13 proteins can be fused deaminase enzymes to edit RNA sequences.^4,12^ Fusion of dCas13 to an array of effectors enables systems that alter^16^ and sense^17^ post-transcriptional modifications such as nucleobase methylation, modulate mRNA splicing, and upregulate translation.^18–20^ In all cases, Cas13 serves as a short guide RNA-programmed domain that binds and stabilizes RNA targets to induce proximity to an effector domain, uniting sequence-specific targeting with functional modules.

Despite the flexibility in function, these systems are limited by off-target activity, unknown interactions with regulatory machinery, and the large size of Cas13.^6,21,22^ Although genome-mining^23,24^ and engineering efforts^25,26^ continue to produce smaller variants, therapeutic applications of Cas-derived complexes encounter challenges with immunogenicity and delivery, as in DNA-targeting contexts. These challenges are compounded for all non-mammalian, protein-based RNA manipulations due to the potential requirement for repeat dosing due to RNA turnover. Reprogramming mammalian cells to perform targeted RNA manipulations offers a transformative option for biotechnological and therapeutic applications. As evidenced by Cas proteins, most biotechnologies for manipulating mammalian cells in a programmable fashion utilize some non-native components, potentially creating homeostatic and immunogenic liabilities while sacrificing the evolutionary optimization of mammalian proteins that could be exploited by a native system.

An existing approach to building programmable RNA effectors with improved immune tolerance and size is to design them in a bottom-up fashion from pieces of mammalian proteins and oligo-RNAs.^27^ Similar to the use of Cas proteins, how the individual pieces of these assemblages interact with the cellular environment, and how the whole construct interfaces with the immune system, remain unknown. Isolation of individual components removes them from their evolutionary context, potentially mitigating advantageous properties and interactions while disrupting components of the endogenous donor protein’s functions or interactomes. As an alternative concept, identifying exogenous reprogramming agents which induce a range of functions by hijacking endogenous pathways could enable conservation of evolutionarily optimized properties while retaining benefits of compatibility with the desired host system. Mammalian cells have the ability to use small RNAs to program the Argonaute (AGO) family of proteins to downregulate a cognate target.^9^ Understanding the RNAi pathway has enabled the usage of engineered small-interfering RNAs (siRNAs) as therapeutic agents that guide AGO2 to a target that is subsequently downregulated.^10^ In principle, human AGO proteins and their native interactors already exhibit many of the advantageous properties of Cas proteins without the need to build a chassis to accomplish target and guide RNA recognition. AGO2-guide RNA complexes can efficiently sample cellular RNAs for targets,^28,29^ and show enhanced binding kinetics and affinity compared to both Cas13^30–32^ and free RNA base-pairing.^28,29^ Despite the advances in harnessing AGO2 for downregulation-based therapeutics and unbiased screening of the transcriptome,^33^ the intrinsic functions of AGO2 are limited. Here, we describe a new concept in RNA manipulation using guide reprogramming of AGO2 to synergize with a fused effector to accomplish targeted manipulations of RNA. Specifically, we develop a C-to-U RNA base editor termed AGO-Led Targeted Editing of RNA (ALTER) which accesses dCas-like function from a native, small RNA-programmable human scaffold.

### Loop-inducing Guide RNAs Enable RNA Editing by AGO2-A3A Fusions

Extensive biological, structural, and biochemical studies of AGO2 activity have elucidated parameters that lead to optimal downregulation.^30,31^ Consequently, these observations provide a map for programming AGO2 with small RNAs towards non-degradative binding. In the context of accessing alternative AGO2-based functions, ideal small RNA guides would pare away unwanted target cleavage and downregulation while retaining useful, evolutionarily optimized properties of RNA-loaded AGO2 such as dramatically increased rates of target binding, high and tunable target affinity,^28,30,31^ the ability to bind despite target secondary structure,^34^ and cellular regulation that promotes efficient target turnover to boost catalytic activity.

The anatomy of siRNAs can be divided into four distinct segments with different effects on target binding and cleavage (Fig. 1a). The first eight bases (g1-g8) comprise the seed region. Other than g1, which is anchored in a pocket of the AGO protein,^35^ complementarity of the siRNA in this region is critical for target binding. Full complementarity between RNA target and siRNA in the central region (g9-g12) marginally reduces target affinity but is required for target cleavage. Base pairing in the supplementary region, g13-g16, is less crucial than in the seed region but increases target affinity. Finally, base pairing in the tail region, g17 to the end of the siRNA, is largely inconsequential for target binding. Therefore, small RNAs with central region mismatches can be used to impart high-affinity binding to a complementary target without inducing target cleavage.^30,31^ Human microRNAs (miRNAs), which do not induce target cleavage but recruit repressive factors, generally operate through the latter binding mode.^36^ However, most miRNAs target the 3’UTR, sometimes with multiple binding sites, in order to effectively mediate downregulation.^37^ Small RNA guides designed to target non-contiguously to unique or non-3’UTR sites in a target transcript could lead to AGO2 binding without potent downregulation. With these parameters, we sought to convert AGO2 to a platform for RNA manipulation beyond downregulation by engineering small RNA guides for a specific function.

**Fig. 1:**
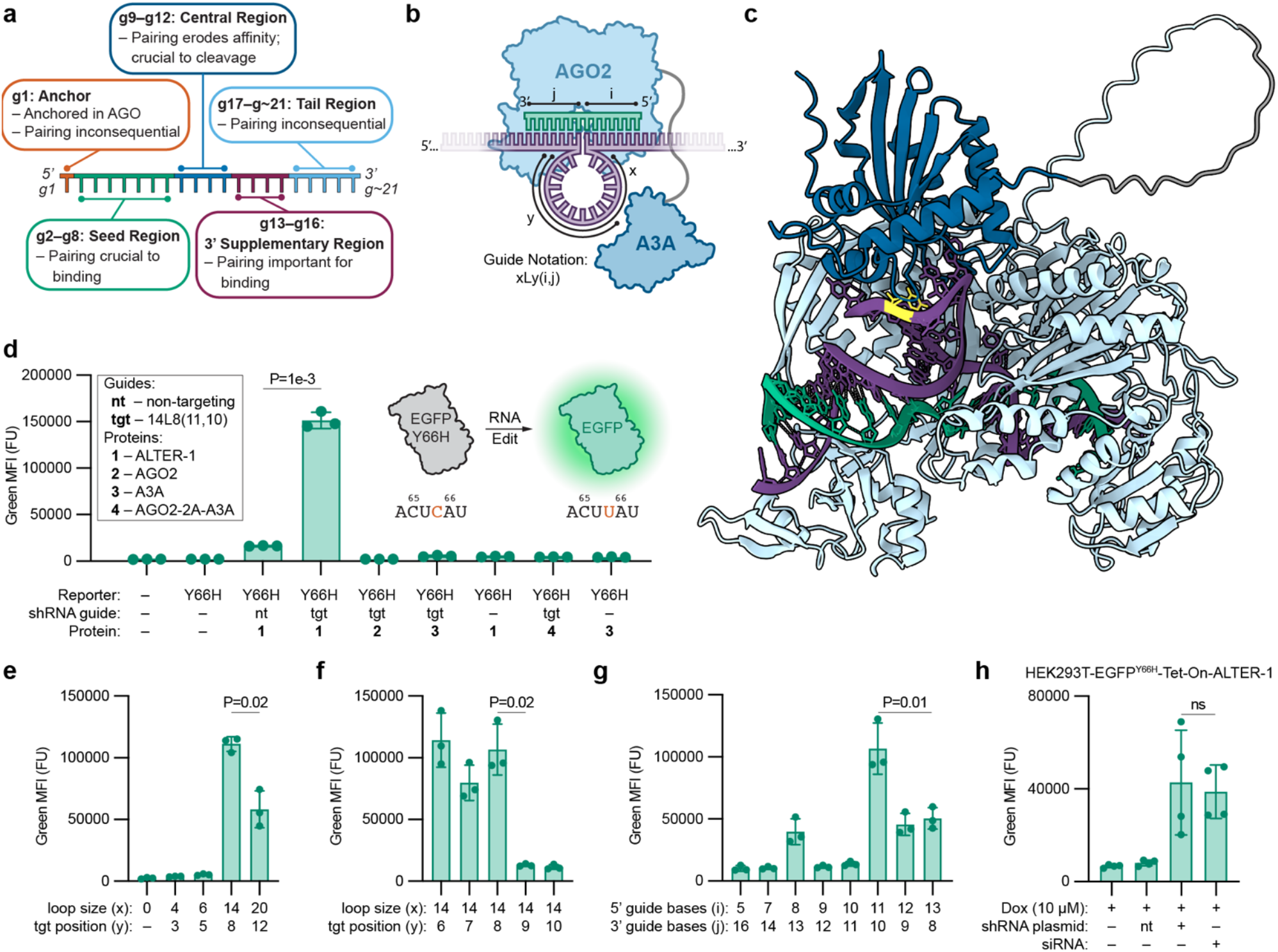
ALTER Design and Guide Optimization. **a**, The functional regions of siRNA and the impact of guide-target base pairing on target affinity and cleavage within each region. **b**, Schematic of ALTER editing. Green: guide RNA, purple: target RNA. Guides are described by the variables: (x) loop size, (y) position of target base in loop (5’ -> 3’, target RNA), (i) 5’ guide bases before the loop, (j) 3’ guide bases after the loop. **c**, AlphaFold3 model of ALTER-1 bound to the EGFP^Y66H^ 14L8(11,10) guide and the targeted portion of the mRNA. Light blue: AGO2, dark blue: A3A, gray: (GGS)_6_ linker, green: guide RNA, purple: target RNA, yellow: target base. **d**, Fluorescence data from HEK293T cells stably expressing EGFP^Y66H^ showing guide-dependent editing and controls that support a loop-induction mechanism. **e**, Tested guides inducing different sized loops (x). **f**, Tested guides inducing 14 nucleotide loops with the target base at different positions within the loop (y). **g**, Tested guides with the 14L8 loop at different positions, with respect to the guide (i, j). **h**, HEK293T cells containing a doxycycline-inducible ALTER-1 used to compare plasmid-encoded shRNAs and exogenous siRNA duplexes as guides. Cells were treated with 25 µM doxycycline. g1–g∼21, guide RNA bases 1–∼21, counting from the 5’ end; AGO2, Argonaute 2; A3A, APOBEC3A; nt, non-targeting guide; tgt, targeting guide; EGFP, enhanced green fluorescent protein; Tet-On, tetracycline-inducible expression system; Dox, doxycycline. All editing readouts were measured by live-cell flow cytometry.

Apolipoprotein B mRNA editing enzyme, catalytic polypeptide (APOBEC) proteins are known to act on single stranded RNAs (ssRNAs) to catalyze a deamination reaction that results in C-to-U sequence editing.^38,39^ Native substrates of APOBEC3A (A3A) position the target C in the 3’ portion of stem-loops.^40^ We envisioned that the central mismatch in an AGO2 guide RNA (gRNA) could be used to induce a ssRNA loop at a target position, creating a favorable editing substrate for an fused A3A protein. This AGO2-A3A fusion (ALTER-1) would therefore be programmed by a loop inducing guide that would lead to editing of the target sequence without substantial downregulation (Fig. 1b). The feasibility of this design was supported by modeling in AlphaFold3^41^ which showed correct folding of the protein fusion components and association of A3A with the unbound portion of the RNA target (Fig. 1c, Extended Data Fig. 1a). C-to-U RNA base editors have the potential to correct pathogenic T-to-C mutations, and while several have been developed, none have been applied therapeutically likely as a function of immunogenicity and efficacy.^4,12,42^

We considered three parameters in gRNA design: (1) the size of the induced ssRNA loop (x), (2) the position of the target C within that induced loop (y), and (3) loop orientation with respect to the gRNA (i and j) (Fig. 1b). We engineered HEK293T cells to express EGFP^Y66H^,^43^ a fluorophore inactivated mutant of EGFP that is rescued by a C-to-U edit (Y66 to H66, Fig. 1d). Transfection with ALTER-1, consisting of wt-AGO2 fused to A3A^Y132D^,^44^ and 14L8(11,10), a guide which induces a 14 nucleotide ssRNA loop between g11 and g12 in which the target C is at position 8,^12^ led to strong induction of fluorescence (Fig. 1d). A non-targeting guide, the absence of either AGO2 or a linked fusion to A3A, or the overexpression of A3A alone did not lead to similar levels of fluorescence. These controls support the proposed mechanism where ALTER-1 binds a target in a guide sequence-defined manner and induces editing. To assess knockdown ability of the 14L8(10,8) guide, we integrated destabilized GFP^45^ into HEK293T cells and transfected non-targeting, fully complementary, and 14L8(10,11) shRNAs. The 14L8(11,10) guide did not lead to downregulation of GFP compared to non-targeting control, demonstrating a lack of activity stemming from endogenous AGO2. Upon transfection of these cells with ALTER-1, we did not observe significant downregulation of GFP by the 14L8(11,10) guide compared a non-targeting guide, but saw marginal downregulation compared to guide without co-expressed ALTER-1 (Extended Data Fig. 1b). Additionally, measurement of stably incorporated EGFP^Y66H^ transcript levels by qPCR did not show a guide-dependent decrease following ALTER transfection (Extended Data Fig. 1c). These results support that loop-inducing guides lead to minimal AGO2-mediated knockdown of the target.

To probe the effects of loop position, size, and orientation, we designed various guides consisting of 21 nucleotides. Alterations in the loop size (x) showed that smaller loops led to an apparent lack of editing, while expansion of the loop size to 20 nucleotides reduced editing efficiency (Fig. 1e, Extended Data Fig. 1d). Shifting placement of the target C within the 14-nucleotide loop (y) showed similar editing when the target was at the 6, 7, or 8 position. Conversely placing the target at position 9 or 10 led to pronounced decreases in fluorescence (Fig. 1f, Extended Data Fig. 1e). Finally, we varied the position of the loop with respect to the gRNA (i and j). These experiments showed that shifting the loop toward the 5’ end of the gRNA sharply reduced editing while shifting to the 3’ end led to a less pronounced decrease (Fig. 1g, Extended Data Fig. 1f). This binding mode, where all bases of the gRNA are complementary to the target, but not contiguous, is atypical for native miRNAs. To examine canonical miRNA binding modes, we designed guides with one or three mismatches in the central region. Both designs decreased editing activity (Extended Data Fig. 1g). Together, these data demonstrate that a novel, loop-inducing binding mode for guide RNAs inside of AGO2 generates favorable substrates for A3A enzymes.

### ALTER Activity Can be Improved by Rational Protein Engineering

AGO2 has been extensively characterized through mutational studies which perturb its RNAi-related functions. Modifications which ablate nuclease activity,^9^ reduce non-cleaving target downregulation,^46^ and minimize the size of the protein through domain truncation^47^ could offer beneficial impacts on neofunction induction (Fig. 2a). While loop-inducing guides bypass proposed requirements for target cleavage, residual slicing of target transcripts could impact ALTER editing efficiency. To test whether complete removal of nuclease activity would lead to greater editing signal, we first investigated whether use of AGO isoforms, AGO1 and AGO3, which do not cleave targets with standard-length shRNAs, would improve ALTER.^48,49^ While ALTER constructs using these isoforms led to guide-dependent editing, the resulting signal was significantly lower than ALTER-1 (Extended Data Fig. 2a). Second, we manipulated AGO2 nuclease activity with several reported mutations (Fig. 2b). D597A and D669A led to decreased editing activity while H634P did not significantly differ in activity from ALTER-1. Conversely, the Q633R mutation led to a pronounced increase in ALTER activity. This mutation has been reported to ablate slicing in certain contexts,^9,49^ but may also increase ALTER activity by increasing affinity to the negatively charged target RNA as modulation of affinity of AGO2 to target mRNA by negative charge induction is a known regulatory mechanism.^50,51^ The double mutant AGO2^F470V, F505V^, located in the AGO2 MID domain (Fig. 2c, left), has been shown to reduce non-slicing downregulation of the target RNA, although the underlying mechanism remains unclear.^46,52–55^ Incorporating this mutation into ALTER resulted in a modest decrease in signal but a dramatic decrease in background signal from the editor with a non-targeting guide, potentially through decreased association with polysomes (Fig. 2c, right). Proximity to mRNAs in polysomes might lead to increases in background and signal which are both removed by AGO2^F470V, F505V^. Together, these efforts identified mutations in AGO2 which could increase editing signal and lower background activity.

**Fig. 2:**
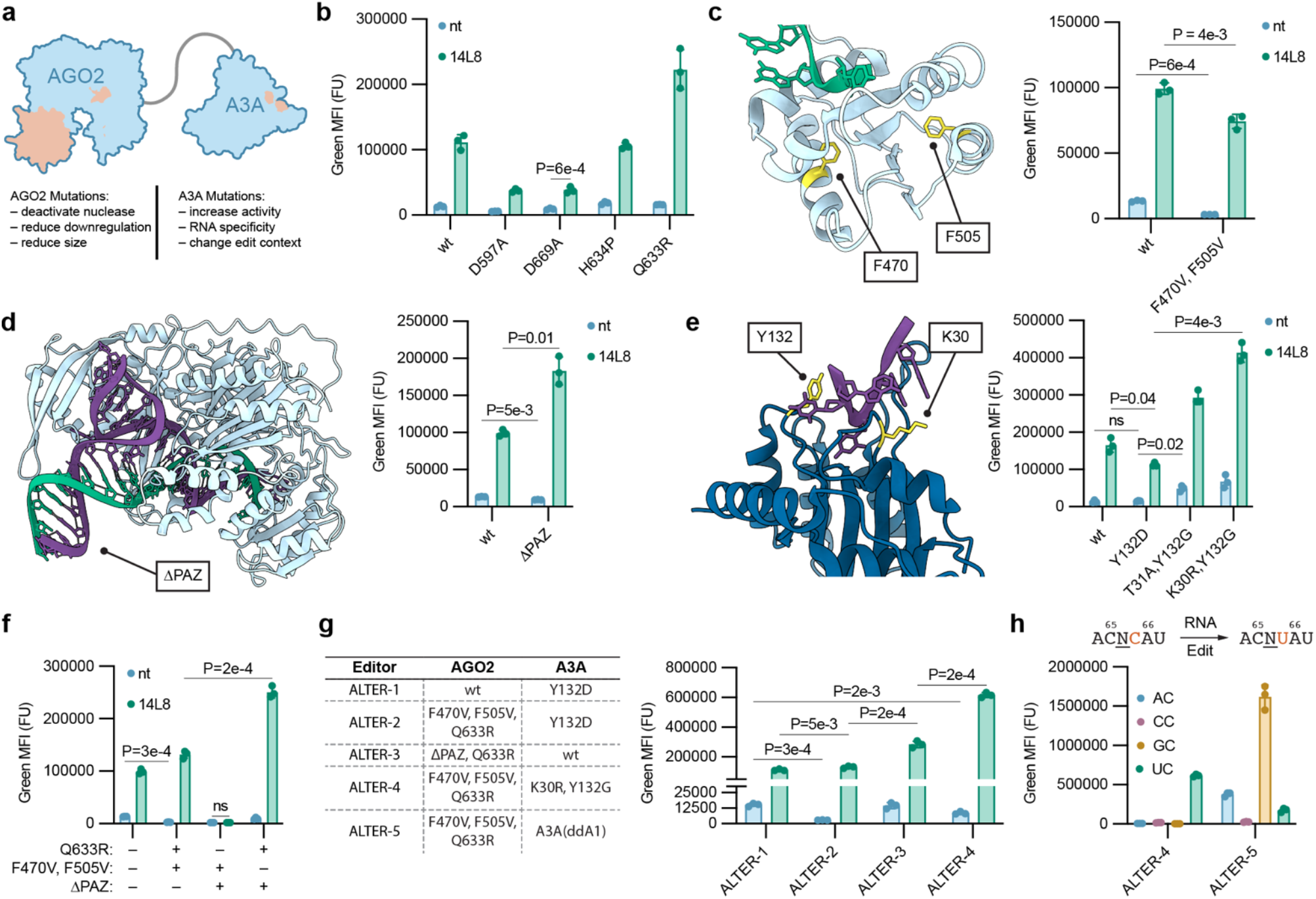
Engineering ALTER with Known Functional Mutations of AGO2 and A3A. **a**, Different properties of AGO2 and A3A that can be modulated with known mutations. Areas with relevant residues highlighted. **b**, Effect of nuclease-inactivating AGO2 mutants in ALTER. **c**, The AGO2 double mutant F470V, F505V in the MID domain reduces non-slicing target downregulation (left, PDB 9CMP).^72^ Effect of AGO2^F470V,F505V^ on activity compared to a nt guide when used in ALTER (right). **d**, AlphaFold3 model of AGO2 lacking the PAZ domain (left). Effect of AGO2^ΔPAZ^ in ALTER. **e**, The effects of different A3A mutants on ALTER (PDB 5SWW).^73^ **f**, Effect of combined AGO2 mutations on ALTER editing efficiency. **g**, Table showing AGO2 and A3A mutations in ALTERs 1–5 (left). Effect of combinatorial AGO2 and A3A mutations on the editing activity of ALTERs 1–4 (right). **h**, Reactivity of chimeric A3A(ddA1) in ALTER-5 in diverse XC substrate contexts. A3A(ddA1), a chimeric APOBEC where the deaminase domain of A1 is grafted into A3A. All editing readouts were measured by live-cell flow cytometry.

The size of protein fusions can be a key determinant of delivery by state-of-the-art vectors such as AAVs, which has instigated numerous efforts in Cas engineering through genome mining^23,24^ or truncation.^25,26^ We reasoned that thorough characterization of AGO2 activity in RNAi studies would enable rational truncation of AGO2. Removal of the 127-amino acid PAZ domain of AGO2 (Fig. 2d, left) has been shown to ablate target cleavage and downregulation while maintaining the ability to load small RNAs and bind to target transcripts.^47^ Incorporating AGO2^ΔPAZ^ into an ALTER editor led to an two-fold increase in signal from edited EGFP(Y66H) and a simultaneous decrease in background signal with the non-targeting guide (Fig. 2d, right). Modeling of AGO2^ΔPAZ^ with AlphaFold3^41^ supported minimal impact on the structure of the other AGO2 domains (Fig. 2d, left).

Further editor optimization was carried out on the A3A portion of ALTER. Mutations of A3A at the residues which interact with the nucleic acid substrate have been shown to modulate activity and specificity, including mitigating activity on DNA. ALTER-1used human A3A^Y132D^, reported to limit off-target activity in both DNA and RNA Cas base editors.^12,44^ Use of A3A^WT^ in ALTER led to a modest increase in editing signal without increased background as gauged with the non-targeting guide. We also tested A3A mutations known to have increased activity on RNA and negligible activity on DNA (Fig. 2e).^56^ While both led to increased background activity, the A3A^K30R,Y132D^ mutant also led to a significant increase in targeted editing.

Having identified mutants in both proteins of ALTER able to modulate on-target and background activity, we investigated the incorporation of multiple mutants into single editors (Fig 4f). AGO2^F470V, F505V, Q633R^-A3A^Y132D^ (ALTER-2), showed minimal signal with the non-targeting guide and higher on-target activity than ALTER-1. ALTER-3, AGO2^ΔPAZ, Q633R^-A3A^WT^, also showed decreased background with higher editing activity than ALTER-2. Constructs containing AGO2^ΔPAZ,F470V, F505V^ mutations reduced editing to barely detectable levels (Extended Data Fig. 2b). Finally, AGO2^F470V,F505V,Q633R^-A3A^K30R,Y132G^ (ALTER-4) produced lower background activity than ALTER-3 with a nearly two-fold increase in editing.

Human A3A is limited by its strict selectivity for UC sequence context. Other human APOBECs have different selectivities, for example APOBEC1 (A1) is requires an AC context for deamination.^39^ However, incorporation of A1 into ALTER did not result in editing (Extended Data Fig. 2c, d). Alternatively, sequence specificity can also be achieved in APOBEC hybrids created with domains from different isoforms. Combining the deaminase domain of A1 with A3A (A3A^ddA1^) has been shown to produce an active GC deaminase.^57^ AGO2^F470V,F505V,Q633R^-A3A^ddA1^ (ALTER-5) showed efficient editing in a GC variant of the EGFP^Y66H^ reporter and significant activity with an AC reporter. ALTER-5 demonstrates that different effector domains can be fused to AGO2 to change function.

### ALTER is a Highly Active Editor of Endogenous RNA Targets

The two classes of Cas-fusion RNA base editors are typified by RNA Editing for Specific C-to-U Exchange (RESCUE)^11^ and C-to-U RNA Editor (CURE).^12^ RESCUE consists of dRanCas13b fused to an engineered adenosine deaminase acting on^50,51^RNA (ADAR) variant which can deaminate cytidine and adenosine (Fig. 3a, right), while CURE is a dPspCas13b-A3A^Y132D^ fusion (Fig. 3a, left). We compared different ALTER editors to RESCUE-S and CURE-N in our EGFP^Y66H^ assay (Fig. 3b). CURE showed minimal EGFP signal, while RESCUE was 3-fold less active than ALTER-4 (Fig. 3b). We confirmed editing of the transcript by sequencing, which showed higher activity for ALTER-4 compared to CURE and RESCUE (Fig. 3c, Extended Data Fig. 3a, b). These activity discrepancies suggest that AGO2-based tools may have complementary or superior activity compared to dCas fusions.

**Fig. 3:**
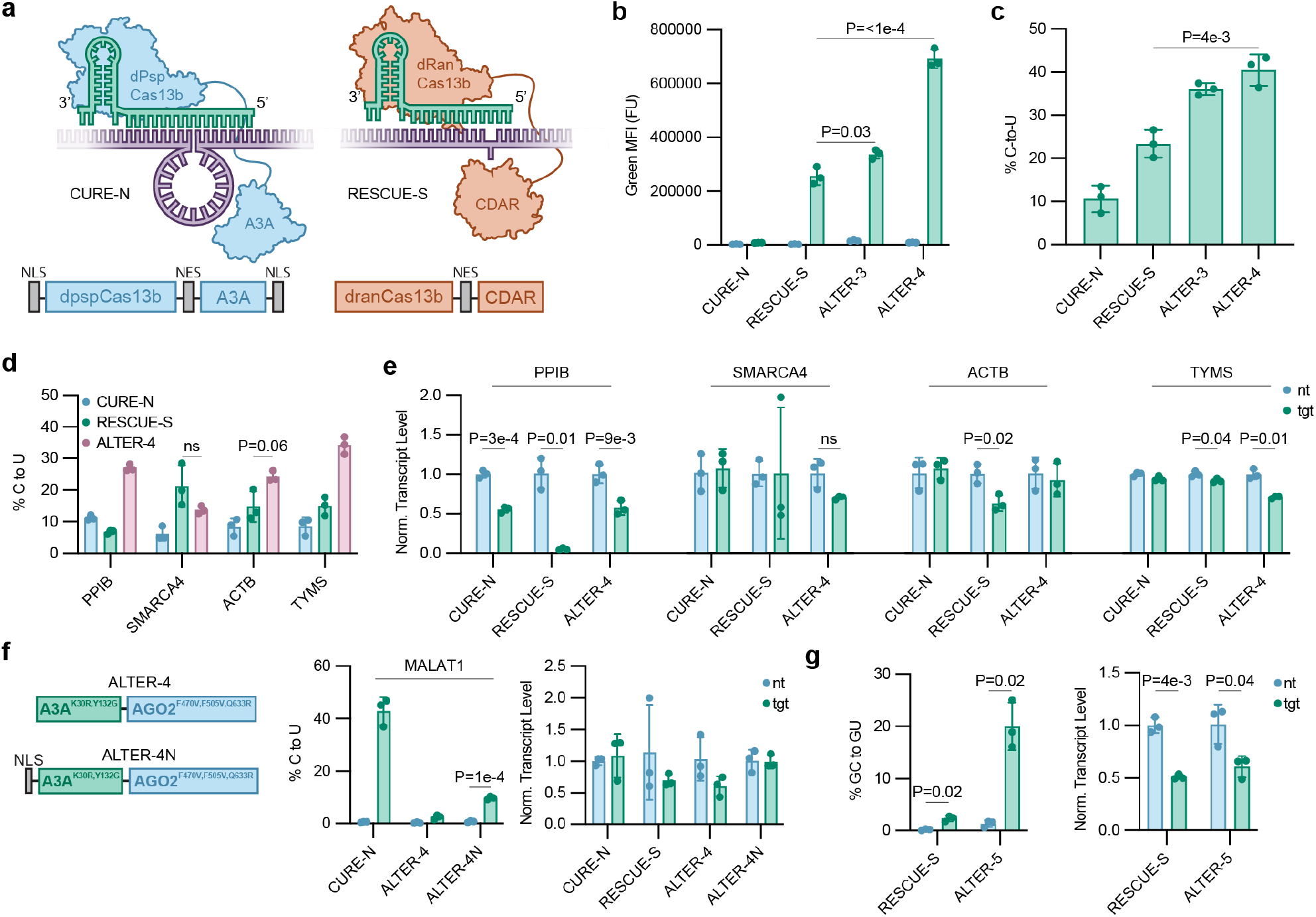
Comparison of ALTER to Cas13-Based RNA Editors and ALTER Editing of Endogenous Transcripts. **a**, Cas13 RNA base editors CURE (left) and RESCUE (right). Schemes showing target binding modes (top) and **editor** components (bottom). **b**, Fluorescence of HEK293T cells stably expressing EGFP^Y66H^ driven by the EF1α promoter treated with different combinations of editors and guides. **c**, Sequencing data showing the percentage of EGFP^Y66H^ transcripts edited at C199 from HEK293T cells co-transfected with a 1:20 mixture of plasmids expressing EGFP^Y66H^ under a PGK promoter and the indicated editor. **d**, Sequencing data showing C-to-U editing on endogenous transcripts in HEK293T cells transfected with different editors. **e**, qPCR data showing relative transcript levels in the endogenous editing experiments. Transcripts were normalized to mean nt level for each editor. **f**, Schemes showing the components of ALTER-4 and ALTER-4N (right). C-to-U editing at in HEK293T cells transfected with plasmids encoding different editors (middle). qPCR data showing relative MALAT1 transcript level (right). **g**, C-to-U editing of PPIB (left) and qPCR data (right) showing relative transcript levels in HEK293T cells transfected with plasmids encoding RESCUE-S or ALTER-5 against a GC target. CURE-N, C-to-U RNA Editor (nuclear localized variant);^12^ RESCUE-S, RNA Editing for Specific C-to-U Exchange (selective variant);^4^ ALTER-4N, ALTER-4 (nuclear localized variant). Target positions by gene, and transcript, and location: PPIB ENST00000300026 C90 (UC), SMARCA4 ENST00000647230 C503, ACTB ENST00000646664 C309, TYMS ENST00000323274 C876, MALAT1 ENST00000534336 C1053, PPIB ENST00000300026 C92 (GC).

We next assessed the target scope of ALTER on a panel of endogenous transcripts. In previous studies PPIB C90 was shown to be more effectively edited by CURE-N than RESCUE-S.^11,12^ We also observed this trend and found that ALTER-4 showed the highest level of editing (Fig. 3d, Extended Data Fig. 3c). SMARCA4 C503 was more effectively edited by RESCUE-S than CURE-N,^11,12^ with ALTER-4 showing comparable editing to RESCUE-S (Fig. 3d, Extended Data Fig. 3d). This suggests that the activity and selectivity of fusion protein editors are likely not solely determined by their effector domains. We also used ALTER-4 to edit ACTB C309 and TYMS C876, two other sites where RESCUE and CURE have been shown to edit effectively.^11,12^ In both cases ALTER-4 led to significantly higher levels of C-to-U editing than either Cas-based editor (Fig. 3d, Extended Data Fig. 3e, f). For all endogenous targets, we also assessed whether ALTER-4 led to knockdown of the targeted transcript and found any reductions in transcript level to be similar to those observed with the nuclease-inactive Cas editors (Fig. 3e). In total, ALTER-4 showed comparable or higher activity than the Cas editors across a panel of endogenous targets without greater target knockdown.

To assess whether ALTER could access nuclear transcripts, we targeted the nuclear lncRNA MALAT1. Initial experiments with ALTER-4 showed minimal editing at MALAT1 C1053 suggesting that ALTER may be largely excluded from the nucleus. However, addition of a nuclear localization sequence (NLS) to the N terminus of ALTER-4 (ALTER-N) led to a pronounced increase in MALAT1 editing without target knockdown (Fig. 3f, Extended Data Fig. 3g). While activity remained lower than CURE-N,^12^ which has been engineered with both viral NLSs and nuclear export sequences (NES), it may be possible to further optimize for nuclear localization or shuttling. ALTER-N did not exhibit differences in editing activity of EGFP^Y66H^ compared to ALTER-4 (Extended Fig. 3h).

We also examined whether ALTER-5 could edit endogenous targets with a GC sequence context. We tested editing at PPIB C92 (which is preceded by G)^11^ and found the editing level to be significantly higher than RESCUE-S and that both editors had similar effects on transcript abundance (Fig. 3g). No CURE variant has been shown to have editing activity in sequence contexts other than UC.^12^

### Assessment of Transcriptome-Wide Off-Target Activity

As ALTER utilizes AGO2 as a core scaffold, transcriptome-wide C-to-U editing may result from loading of endogenous miRNAs.^58^ To quantify transcriptome-wide editing activity, we compared mRNAseq data of cells treated with PPIB-targeted ALTER-4 to a nanoLuc transfection control. We observed only 375 high-confidence off-target edits, which was two-fold lower than the corresponding number reported for RESCUE-S, and five-fold lower than that reported for CURE-N (Fig. 4a).^12^ We further analyzed the RNA sequence context of each off-target (position 0) and found a consensus sequence from −4 to +1 which matches that previously identified for A3A activity (Fig. 4b).^59^ By modeling the secondary structure in the context window we also observed a preference for a four base loop with a six base pair stem, which agrees with the known A3A preference for stem-loop substrates (Fig. 4c, d).^59^ Alignment of the ALTER-4 guide with a broader sequence context did not reveal an increased frequency of seed-binding sites, which would be expected if these off-targets resulted from binding of the AGO2 portion (Extended Data Fig. 5a-b).^10^ We also analyzed differential mRNA expression across editors and found that ALTER results in higher numbers of differentially expressed genes (DEGs) compared to those reported for Cas-based editors (Ext. Data Fig. 4a). However, Gene Set Enrichment Analysis (GSEA)^60^ of all assayed genes did not reveal any significantly dysregulated pathways (Extended Data Fig. 4b, c), and gene ontology enrichment analysis of DEGs did not reveal any bias in the impact of ALTER (Extended Data Fig. 4d-g). We further assessed whether known miRNA target transcripts^61^ were disproportionately represented in the DEGs and observed the frequency to be unchanged between DEGs and all assayed transcripts (Fig. 4e). These data support that ALTER does not disproportionately impact the miRNA regulome. Additionally, we did not observe pronounced enrichment of identified C-to-U off-targets among DEGs caused by ALTER-4 (Fig. 4f). Overall, these results suggest that ALTER C-to-U off-target edits occur predominantly in substrates favorable for A3A activity and do not result from increased ALTER association or cross-loading of ALTER with endogenous miRNAs. This conclusion is further bolstered by the observation of similar sequence-structure consensus for all A3A-based editors but not for RESCUE-S (Extended Data Fig. 5c-j). Further, ALTER results in significantly fewer C-to-U off targets than A3A overexpression. Together these results demonstrate that ALTER off-targets result primarily from increased expression of the fused A3A, but that A3A off-targets are reduced by fusion to AGO2.

**Fig. 4:**
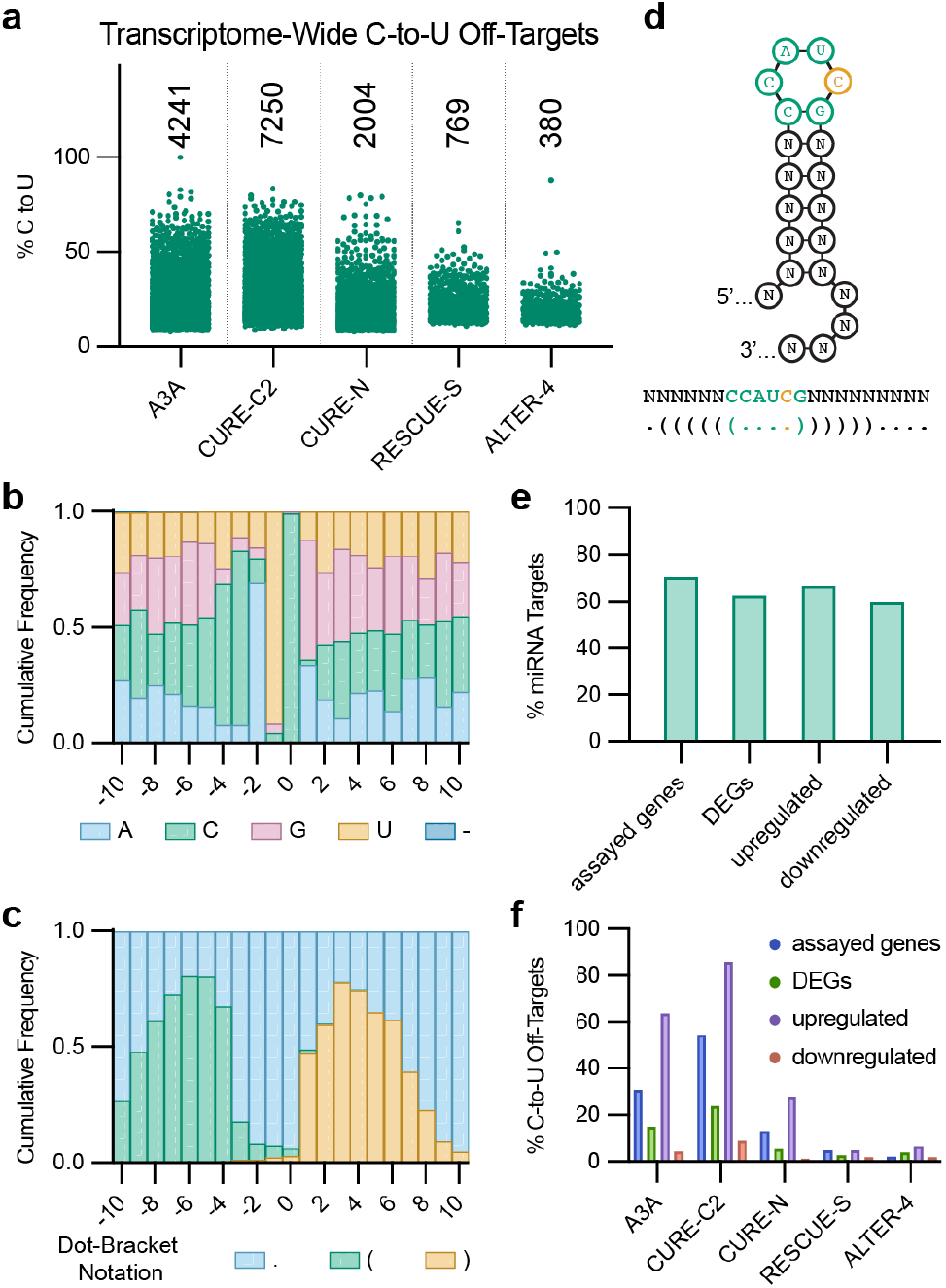
Quantification of Transcriptome-Wide Effects of ALTER. **a**, Quantified C-to-U off targets identified in RNAseq data from HEK293T cells transfected with plasmids encoding each editor. RNAseq data for A3A overexpression, CURE editors, and RESCUE-S obtained from BioProject PRJNA635732.^12^ **b**, Nucleotide frequency in a 21 base context window around each ALTER-4 C-to-U off-target indicating a CCAUCG consensus sequence in the –4 – +1 window. **c**, Frequency of dot-bracket notation characters in dot-bracket encoding of the minimum free energy secondary structure for each 21 base context window. **d**, Scheme of the consensus sequence and structural features of the ALTER-4 C-to-U off target sites (top). Consensus sequence and dot-bracket notation of the consensus structure (bottom). **e**, Frequency of endogenous miRNA targets in all genes with sufficient transcript expression for analysis and different DEG subsets for ALTER-4. **f**, Frequency of C-to-U off targets in all genes with sufficient transcript expression for analysis and in DEG subsets for each editor. CURE-C2, C-to-U RNA Editor (active variant);^12^ DEG, differentially expressed gene; miRNA, microRNA.

### Machine Learning-Assisted Deimmunization of ALTER

As ALTER is based on intact human proteins, we assessed whether sequence engineering could further reduce any predicted immunogenicity without compromising activity. We focused our immunogenicity assessment on MHC class II presentation as activation of cytotoxic T cells requires CD4^+^ T-cell assistance, and prior work indicates that evasion of MHC II–restricted antigen presentation is critical for long-term persistence of engineered or therapeutic human cells. We analyzed ALTER constructs for predicted non-native peptide presentation using MHC Analyzer with Recurrent Integrated Architecture (MARIA), a deep neural network trained on mass-spectrometry-derived antigen data and conventional binding measurements to predict MHC class II presentation.^62^ MARIA has previously been applied to reduce predicted immunogenicity of engineered human proteins.^63^ We computationally generated all 15-mer peptide tiles across a given ALTER variant that contained either an amino acid that differed from the native protein or a junction between two native proteins. These peptides were screened with MARIA and considered likely to be presented if they exceeded the 63rd percentile of total peptide scores, representing a false omission rate (FOR) of 0.001. Similar analyses were performed for CURE-N and RESCUE-S for comparison. As expected, the Cas13-based editors exhibited substantially more likely-presented peptides compared to ALTER editors (Fig. 5a, Extended Data Fig. 6a).

**Fig. 5:**
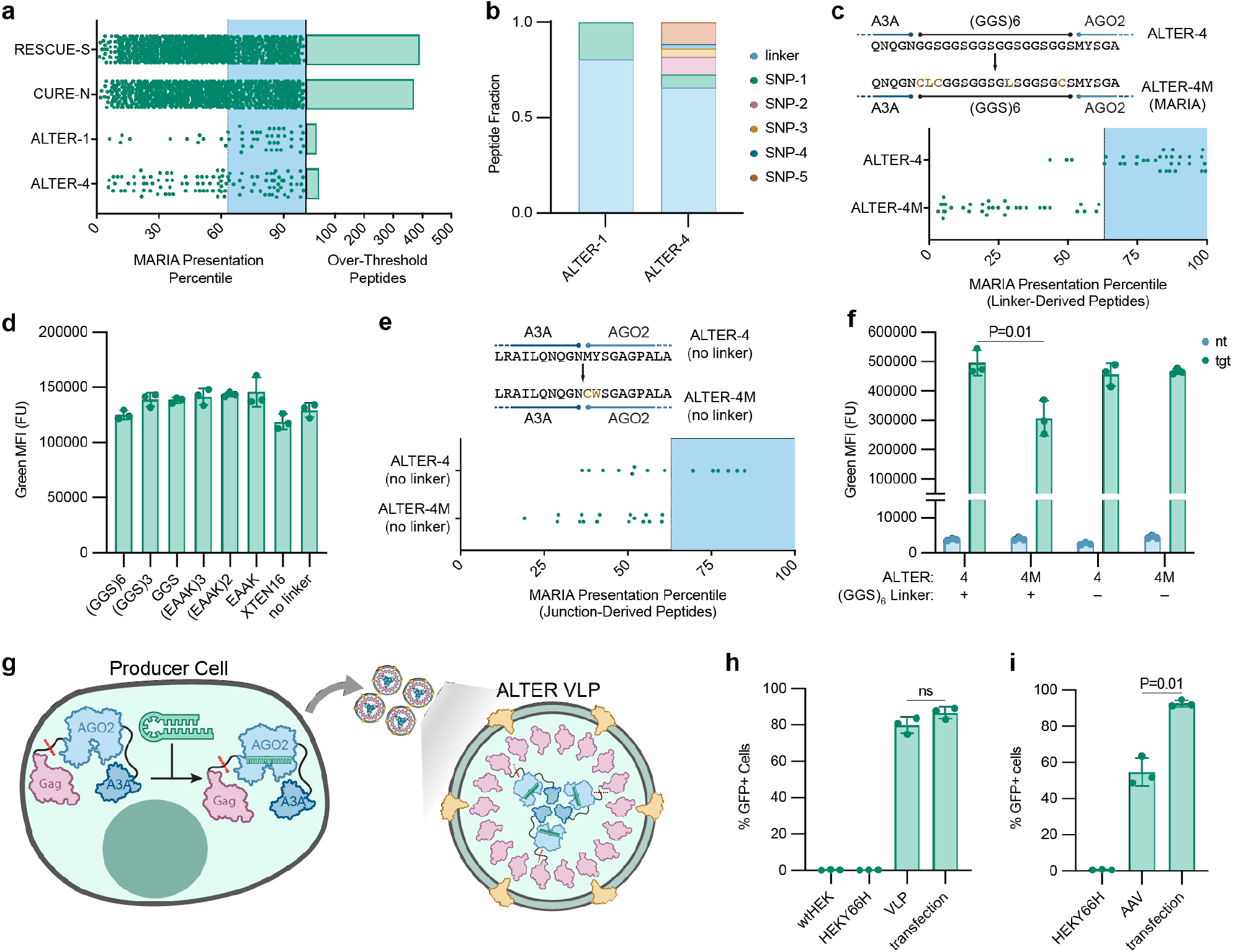
Computational Assessment of Immunogenic Liability and Delivery Modes for ALTER. **a**, MARIA presentation percentile of non-human peptides resulting from different editors and counts of number of peptides exceeding the confidence threshold. **b**, Cumulative fraction of over-threshold peptides from ALTER-1 and ALTER-4 grouped by location of origin. **c**, Peptide sequences showing the de-immunizing mutations in ALTER-4M (top), and MARIA presentation percentile for linker-derived peptides from ALTER-4 and ALTER-4M (bottom). **d**, Editing activity in HEK293T cells stably expressing EGFP^Y66H^ transfected with plasmids encoding ALTER-1 with different linkers and the 14L8(11,10) guide. **e**, MARIA presentation percentile for junction-derived peptides from ALTER-4 (no linker) and ALTER-4M (no linker, bottom). Peptide sequences showing the de-immunizing mutations in ALTER-4M (no linker, bottom). **f**, Editing activity in HEK293T cells stably expressing EGFP^Y66H^ transfected with plasmids encoding ALTER-4, ALTER-4 (no linker), and their de-immunized variants. **g**, VLPs can be produced with loaded ALTER RNPs. Sites where the MMLV-Gag-ALTER linker is cleaved by the viral protease are indicated with dashed red lines. **h**, Percentage of cells positive for fluorescence after treatment with ALTER-4-containing VLPs compared to transfection with an ALTER-4 encoding plasmid. **i**, Percentage of cells positive for green fluorescence after treatment with an ALTER-4-encoding AAV or transfection with the corresponding plasmid. MARIA, MHC Analyzer with Recurrent Integrated Architecture; XTEN16, SGSETPGTSESATPES; ALTER-4M, ALTER-4 with de-immunizing mutations; VLP, virus-like particle; AAV, adeno-associated virus.

For ALTER-1 and ALTER-4, the majority of peptides above the MARIA threshold originated from the (GGS)_6_ linker (Figure 5b). As mutations within linker peptides are less likely to reduce activity than mutations within functional domains, we focused on reducing this major source of predicted immunogenicity. We found that five-point mutations in the linker region (G200C, G201L, S202C, G210L, G216C) eliminated predicted non-native peptide presentation associated with the linker region in ALTER-4 (Fig. 5c). Having identified the linker as a potential focal point of immunogenic liability, we tested the use of a variety of linkers in ALTER. We assayed different lengths of flexible (GGS) linkers, helical (EAAAK) linkers,^64^ the XTEN-16 linker,^65^ and a construct where A3A is fused to AGO2 without a linker (Extended Data Fig. 6e). In our EGFP^Y66H^ assay we found all constructs gave comparable activity (Fig. 5d). This is likely due to the first 35 residues of the AGO2 Beam acting as a flexible linker due to a lack of defined structure (Extended Data Fig. 6e).^66^ We identified two minimally disruptive mutations to AGO2 (M1C, Y2W) that eliminated predicted junction-associated non-native MHC II peptide presentation from no-linker ALTER-4 (Fig. 5e, Extended Data Fig. 6b). The de-immunized ALTER-4 analog, ALTER-4M, resulted in only slightly decreased GFP editing, while the no-linker variants were both indistinguishable in activity from the original ALTER-4 (Fig. 5f). These results indicate that ALTER constructs can be systematically optimized to reduce predicted immunogenicity and reveal an inherent compatibility of AGO2 with effector domains, requiring minimal artificial linker engineering.

### ALTER Can Be Delivered as an RNP or Gene

Translation of several Cas-based technologies has been challenged by their size and integration into state-of-the-art enveloped delivery technologies. While advances in protein engineering to identify smaller constructs and optimization of intact ribonucleoprotein complex delivery have enabled breakthroughs, nucleic acid editing is still limited by delivery capabilities. Virus-like particles (VLPs) can deliver recombinant proteins to cells by linking them to viral capsid proteins with cleavable linkers.^67^ We formulated VLPs so that ALTER would be loaded with guide and packaged into the VLP within producing cells, eliminating the need for guide expression in target cells (Fig. 5g).^68^ The resultant VLPs were able to deliver active ALTER RNPs to cells expressing only the stably integrated EGFP^Y66H^ reporter, resulting in a comparable frequency of editing to that observed with transient transfection (Fig. 5h), though with lower signal magnitude (Extended Data Fig. 6c).

Adenovirus-Associated Vectors (AAVs) enable the delivery of single constructs within a 5 kilobase size limit and are clinically validated vectors for multiple gene therapies.^69^ We designed AAVs that incorporated both the ALTER protein-fusion and guide into a single viral vector.^70^ Treatment of EGFP^Y66H^ reporter cells with these AAVs showed effective editing with only a modest reduction in cell turn-on compared to transfection (Fig. 5i), though with lower signal magnitude (Extended Data Fig. 6d). Together, these results suggest that current technologies can deliver functional ALTER as both an RNP complex and encoded within a gene, supporting multiple potential avenues for therapeutic application of AGO2-based RNA-manipulating systems.

## Discussion

Cas and ISC proteins have led to programmable RNA effectors with a variety of functions^12,15–20,24^ but are not evolutionarily optimized for function in human cells, presenting implications for immune responses and deleterious intracellular effects.^6,21,22^ These shortcomings could be addressed by building similar programmable effectors from intact human proteins. By fusing AGO2 to a human RNA effector protein and engineering the AGO2 guide RNA to prevent target slicing,^30,31^ AGO2 can be redirected from RNAi to enable the function of the fused domain. As a proof-of-concept, in ALTER, a loop-inducing guide that does not lead to substantial target downregulation is used to force the target transcript into an optimal conformation for editing by a fused deaminase. C-to-U editing has the potential to correct a number of pathologies,^42^ and along with Cas-based systems, has been explored with PUF protein scaffolds^44^ or engineered long RNA oligo-guided ADAR2 mutants.^71^ In contrast to these approaches, ALTER features reduced immunogenic liability due to lack of multiple fusions of modified proteins and requiring only short RNAs as guides,^63^ improved ability to sample for targets,^28^ enhanced target binding kinetics and affinity,^29–31^ undetectable activity of A3A^K30R,Y132G^ on DNA,^56^ and a lack of A-to-I off-targets compared to engineered ADAR-based technologies.^11,12,71^ ALTER does not require nuclear entry to achieve effective editing, further minimizing risk to DNA. Engineering of ALTER benefitted from the wealth of studies carried out on human RNAi, enabling rapid identification of modifications that improved ALTER editing, reduced nonspecific background, and reduced construct size. These results suggest that, with respect to both AGO2 modification and guide design, different properties are optimal. This observation may enable insights into AGO2 activity and regulation that might not be gained from studying RNAi.

Some functions that can be effectively mediated with Cas-based effectors may not be accessible using only human proteins, and the use of ALTER may result in a higher rate of DEGs than Cas-based editors. Initiating optimization protein engineering efforts from human starting points could bypass these challenges, and DEGs may be further suppressed by discovering mutations that are deleterious to RNAi and uniquely valuable to building AGO2-based effectors such as those used in ALTER. Our analysis of immunogenic liability with MARIA^62^ supports a substantially lower rate of MHC II presented exogenous peptides with modified AGO2 compared to Cas. Further efforts to expand AGO2-based effectors will continue to benefit from the existing RNAi knowledge to inform new engineering routes. While a subset of the AGO interactome might pose challenges for biotechnological applications, we have demonstrated that targeted mutations may abolish undesired activities while retaining productive features. The ability to achieve this balance for a given application is likely to be context and potentially target-dependent.

ALTER can be delivered as an RNP or as a gene using enveloped delivery vehicles. Yet, delivery with VLP and AAV vectors still presents immunogenic risks.^69^ However, continued advances in deimmunization of these and other protein-delivery technologies may change the primary immunogenic risk from the delivery vector to the protein cargo itself. Both this work and studies of Cas-based RNA effectors suggest that determinants of cellular activity are target-dependent, multi-factorial, and difficult to predict. Indeed, different and sometimes complementary activities between ALTER and Cas base editors were observed. Future RNA-manipulation applications are likely to benefit from an increased array of options that can be surveyed to induce the desired effect. Reprogramming AGO2 expands the scope of modular targeting domains, guided by small RNAs, that can be used in RNA manipulating technologies and shows that they can be built entirely out of existing human proteins.

## Acknowledgements

This work was supported by an NIH F32 fellowship to R.F.L, a Stanford Bio-X Fellowship to S.M.V.H., and a Burroughs Wellcome Fund CASI award to S.M.B.

## Author Contributions

R.F.L and S.M.B. conceived of the project. R.F.L., S.M.V.H, and C. M. conducted experiments and analyzed all data. R.F.L and S.M.B. wrote the manuscript. S.M.B. provided supervision.

## Extended Data Figures

**Extended Data Fig 1:**
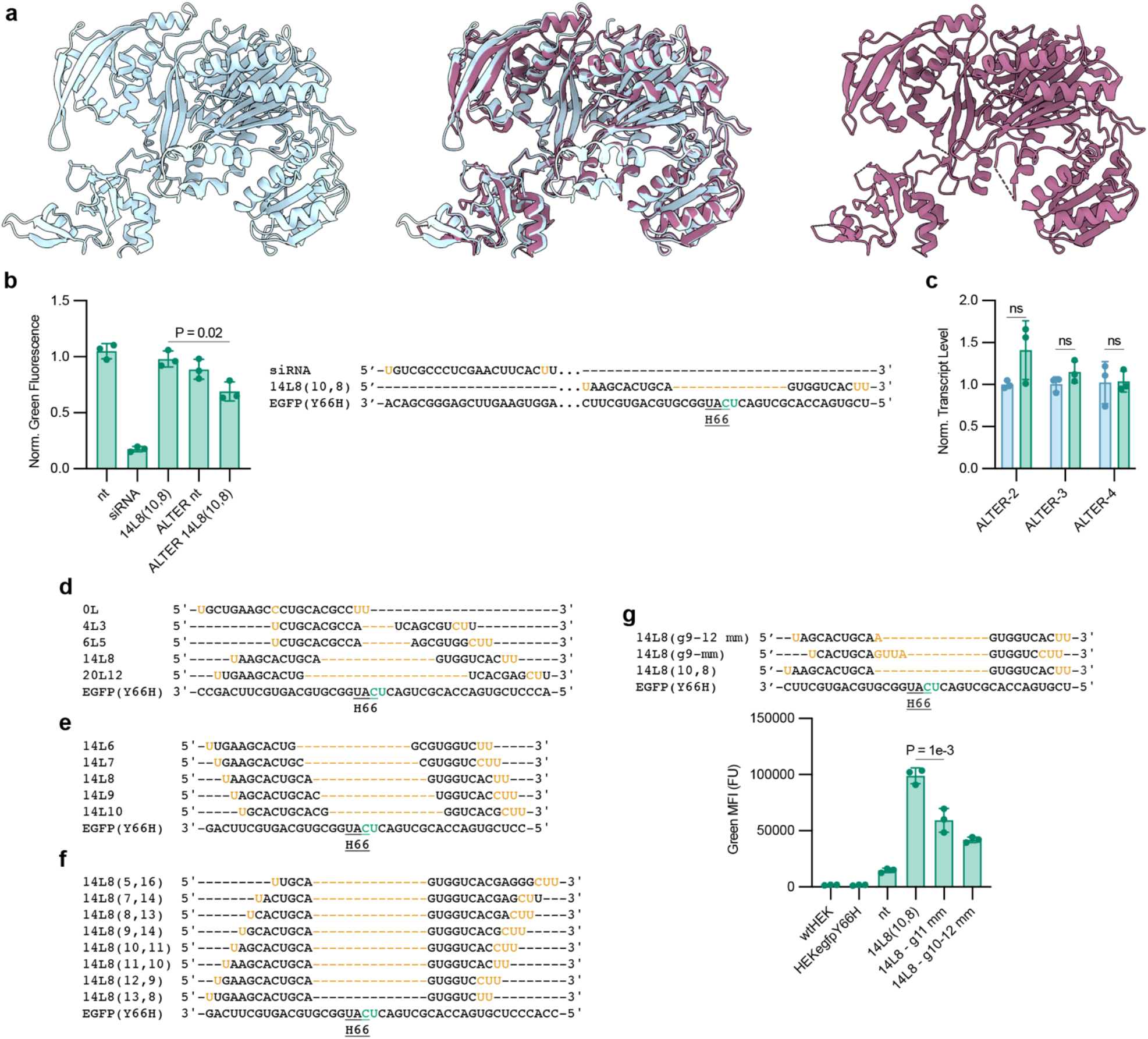
Structural Modeling of AGO2 and Additional Guide Optimization. **a**, The AGO2 domain of ALTER modeled with AlphaFold3 (left) overlaid with known AGO2 crystal structures (right, PDB 6N4O).^36^ **b**, Guide sequences and expected base pairing to a ddGFP target (top). Fluorescence data from HEK293T cells stably expressing ddGFP driven by an EF1α promoter transfected with plasmids encoding an shRNA or shRNA+ALTER-1 (normalized to untransfected cells, bottom). **c**, qPCR of EGFP^Y66H^ transcripts in cells stably expressing the reporter driven by the PGK promoter after transfections with different ALTER-guide combinations. Guide sequences and expected base-pairing to target (top) and resulting fluorescence data (bottom). Guide sequences and expected base pairing for the guides used in: **d**, Fig. 1e. **e**, Fig. 1f. **f**, Fig. 1g. **g**, Testing guide sequences featuring additional mismatches to the EGFP^Y66H^ target. Guide sequences and expected base-pairing to target (top) and resulting fluorescence data (bottom).

**Extended Data Fig 2:**
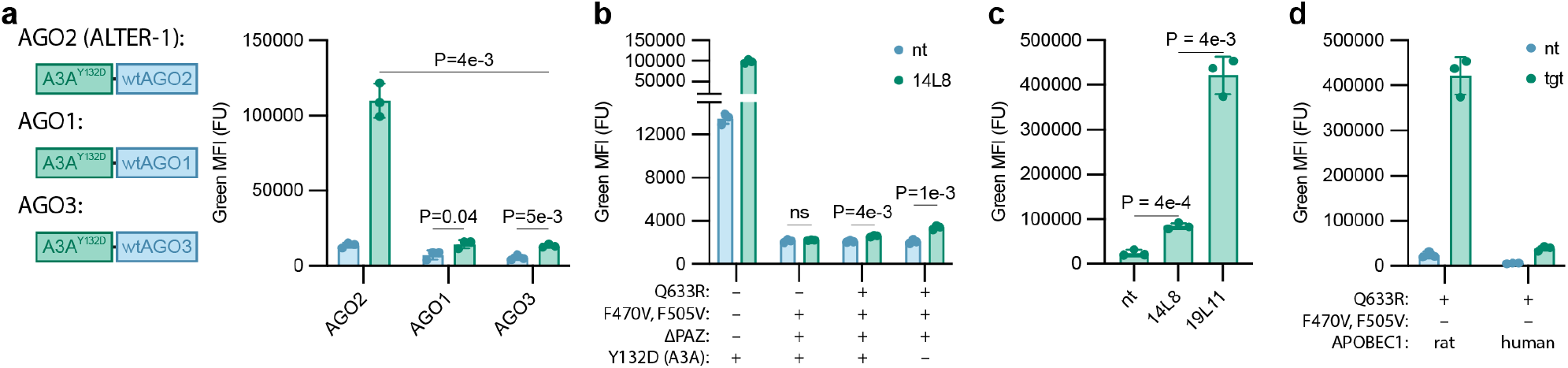
Expander ALTER Modifications. **a**, EGFP^Y66H^ editing activity of ALTER editors using AGO1 or AGO3 as the targeting system. **b**, ALTER editors with both F470V, F505V and ^Δ^PAZ modifications. **c**, Effect of rat A1 in ALTER on an AC EGFP^Y66H^ reporter construct using a19L11(11,10) guide. **d**, Effect of human A1 in ALTER editing of the AC EGFP^Y66H^ reporter construct.

**Extended Data Fig 3:**
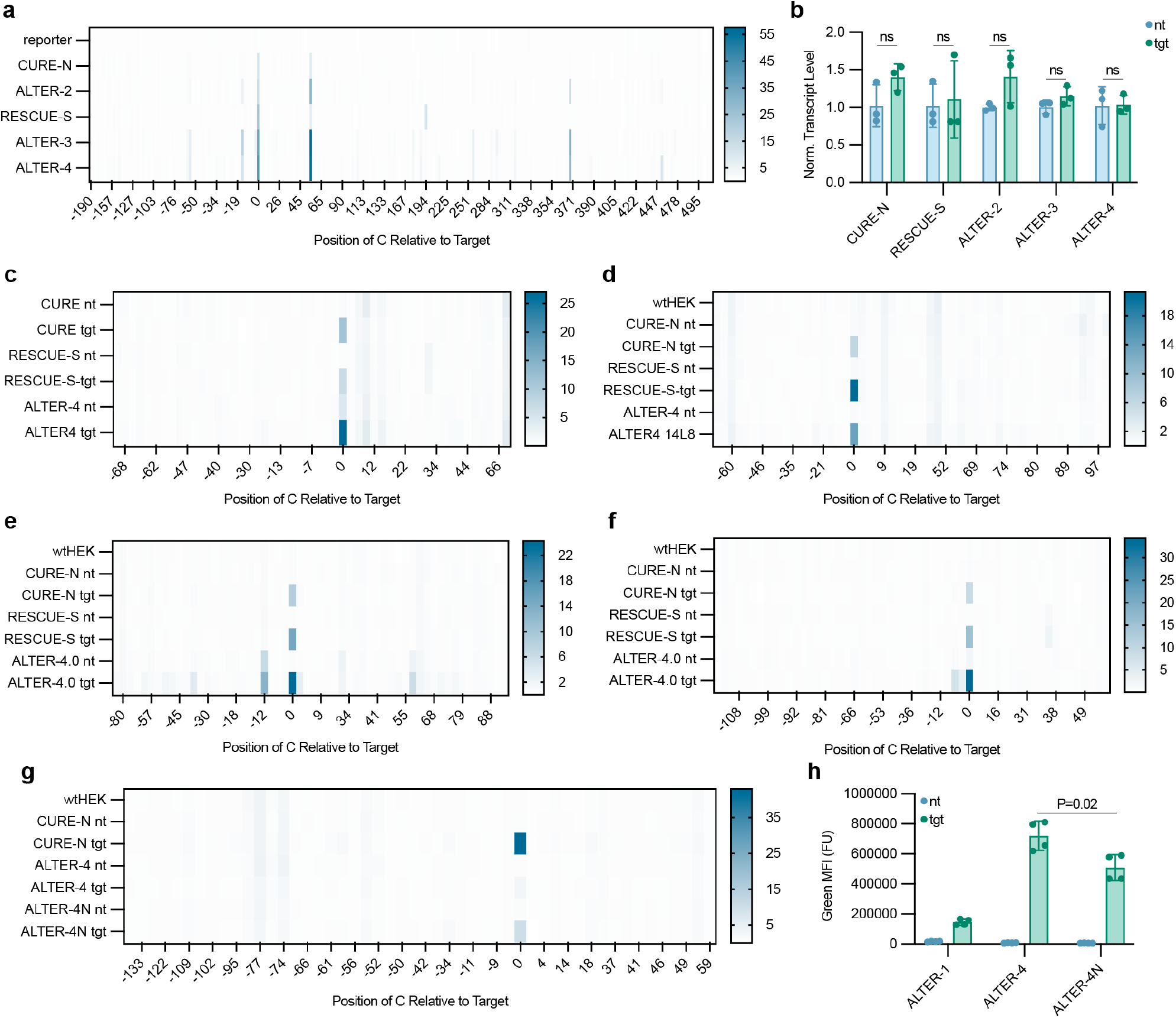
Editing Activity and Off-Targets on Endogenous Targets. **a**, Heatmaps showing C-to-U editing activity at all C nucleotides in EGFP^Y66H^. **b**, qPCR data for PGK-driven EGFP^Y66H^ stably incorporated into HEK293T cells and transfected with plasmids encoding different editors. Transcripts were normalized to mean nt level for each editor. **c–g**, Heatmaps showing C-to-U editing activity at all C nucleotides in the sequencing amplicons used for each endogenous target: **c**, PPIB ENST00000300026 C90. **d**, SMARCA4 ENST00000647230 C503. **e**, ACTB ENST00000646664 C309. **f**, TYMS ENST00000323274 C876. **g**, MALAT1 ENST00000534336 C1053. C positions are given relative to the target C, which is at 0. **h**, Fluorescence data from EGFP^Y66H^ editing with ALTER-4N.

**Extended Data Fig 4:**
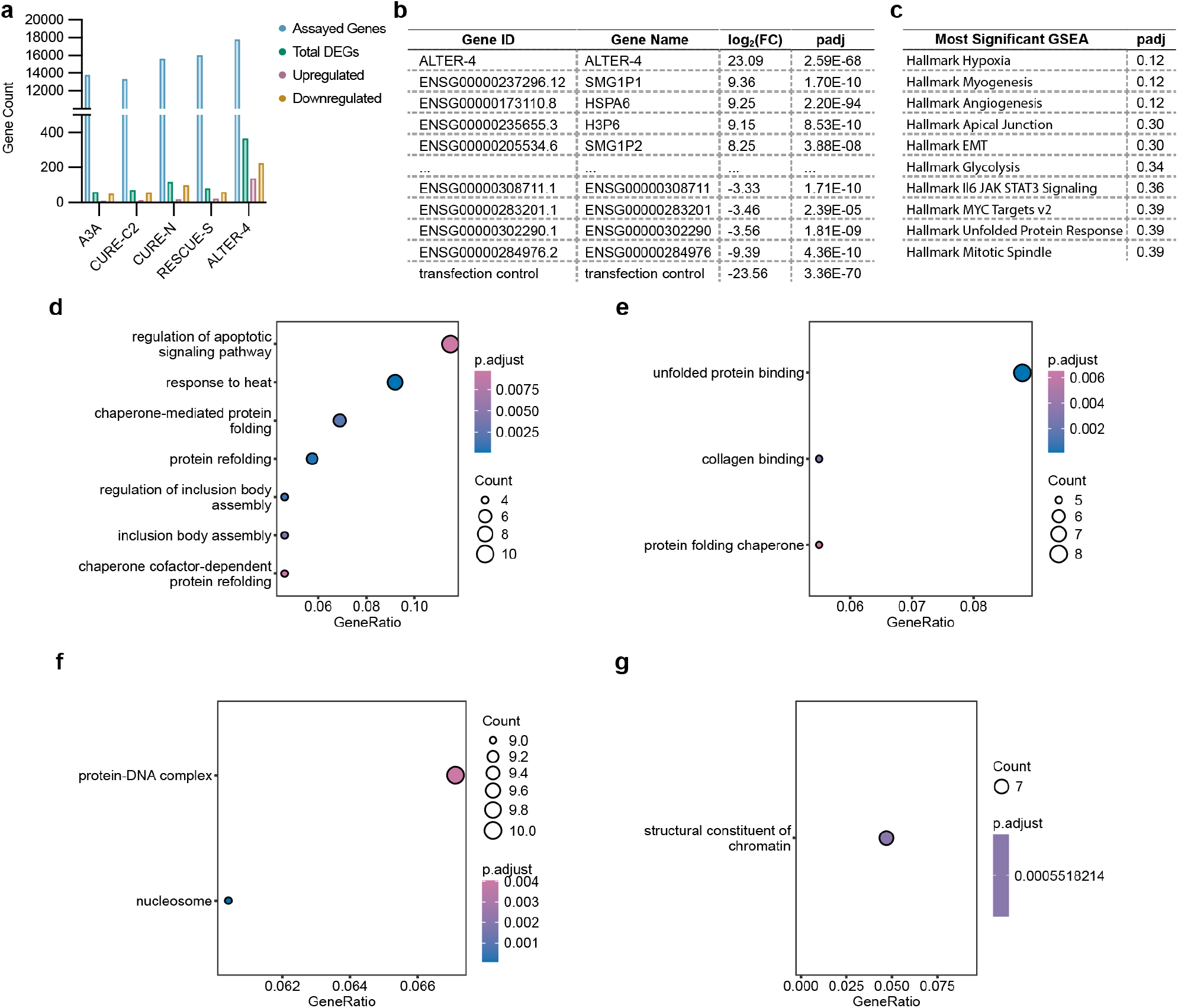
ALTER-4 Transcriptome-Wide DEGs. **a**, Counts of assayed genes and different DEG subsets from RNAseq data for different editors. RNAseq data for A3A overexpression, CURE editors, and RESCUE-S from BioProject PRJNA635732.^12^ **b**, List of assayed genes ranked by their log_2_(Fold Change), compared to the nanoLuc transfection control, in the ALTER-4 RNAseq data. **c**, GSEA results for ranked ALTER-4 log_2_(FC) compared to transfection control.^74^ Dot plots of gene ontology clustering analysis: **d**, upregulated DEGs, biological process GO terms. **e**, upregulated DEGs, molecular function GO terms. **f**, downregulated DEGs, cell component GO terms. **g**, downregulated DEGs, molecular function GO terms. For analysis of upregulated DEGs by cell component GO terms there were no hits. For analysis of downregulated DEGs by biological process GO terms there were no significant hits.

**Extended Data Fig 5:**
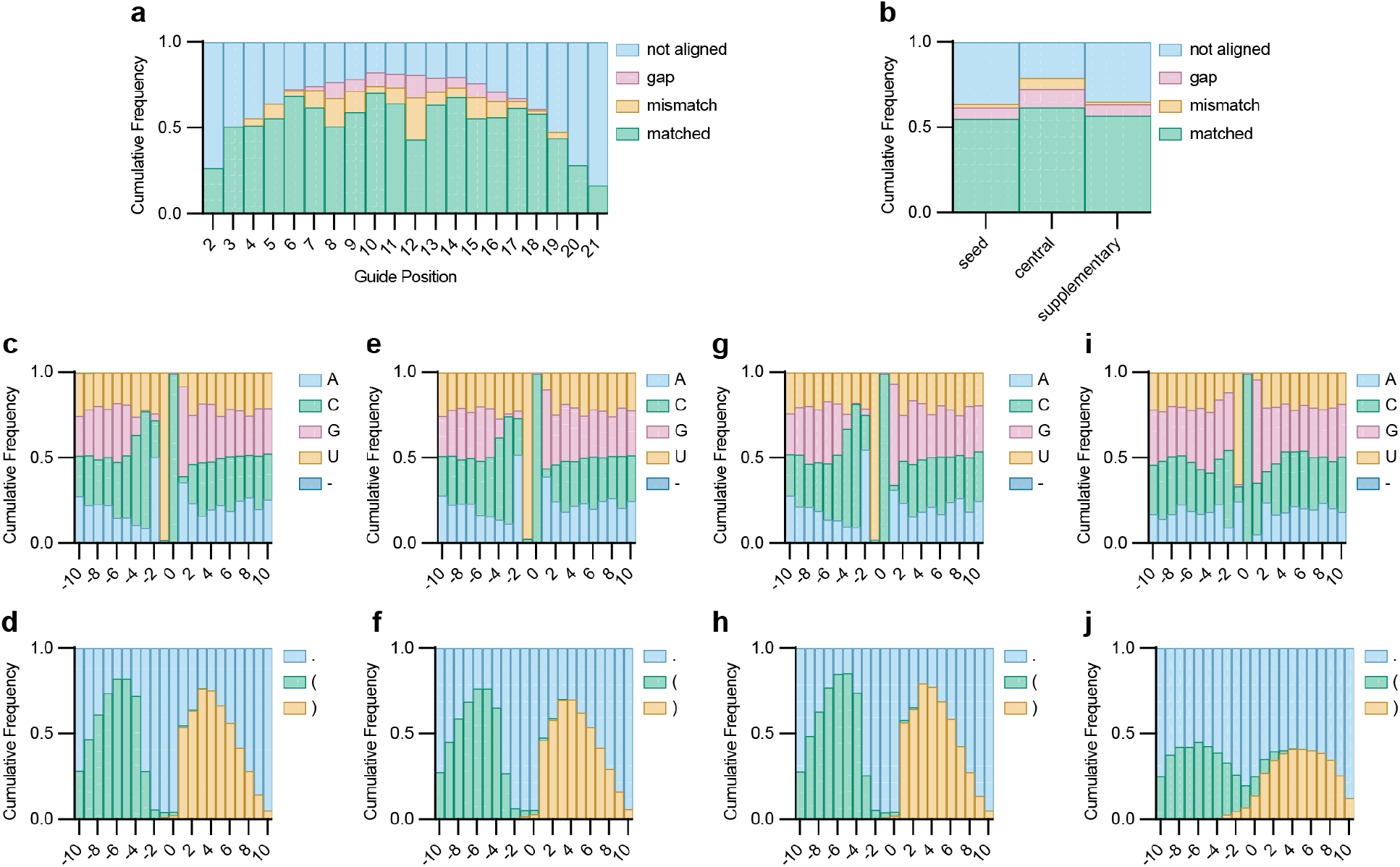
ALTER Guide Alignment Analysis of Regions Surrounding C-to-U Off-Targets and Sequence-Structure Consensus Analysis for C-to-U Off-Targets in Other Editors. **a**, Frequency of alignment outcome at each position in the ALTER guide following basic alignment to a 51 base window centered on each identified off-target. **b**, Alignment outcome data grouped by guide region. Nucleotide (top) and structural element in the lowest MFE secondary structure (bottom) frequencies for each position in 21 base context windows centered around each off target for: **c, d**, A3A overexpression. **e, f**, CURE-C2. **g, h**, CURE-N. **i, j**, RESCUE-S. RNAseq data for **c**-**h** from BioProject PRJNA635732.^12^

**Extended Data Fig 6:**
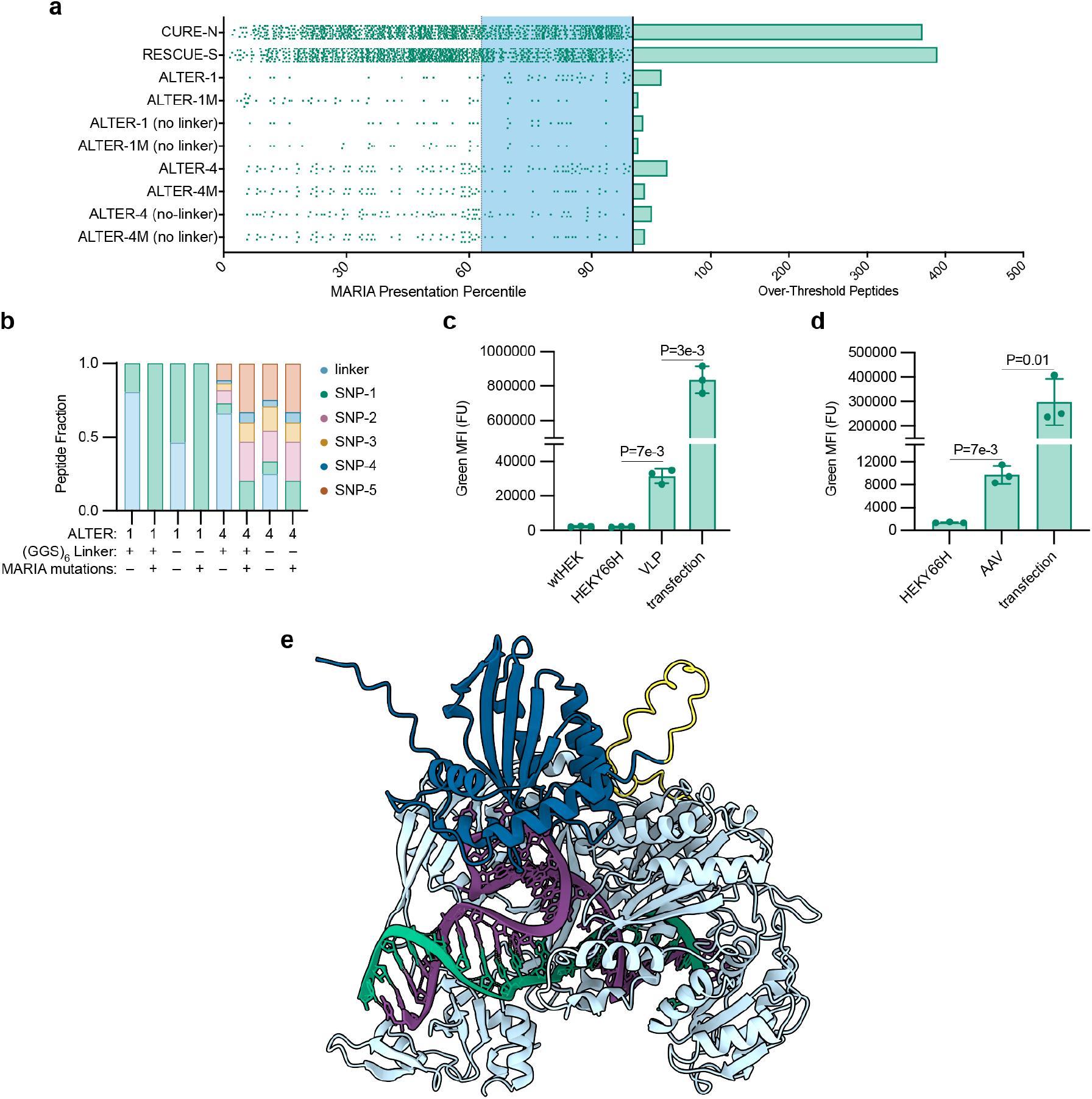
MARIA Analysis of Different RNA Editors and Fluorescence Intensity After ALTER Delivery Via Enveloped Particle. **a**, MARIA presentation percentiles for all non-human 15-mer peptides resulting from each RNA editor discussed (left). Counts of over-threshold peptides for each editor discussed (right). **b**, Fraction of over-threshold peptides resulting from each non-human modification in each ALTER variant along with their de-immunized analogs. **c**, Fluorescence of HEK293T cells expressing EGFP^Y66H^ and treated with ALTER-4-loaded VLPs. **d**, Fluorescence of HEK293T cells expressing EGFP^Y66H^ and treated with AAVs encoding ALTER-4. **e**, ALTER-1 lacking a linker between AGO2 and A3A modeled with AlphaFold3.^41^

## Methods

### Cell Lines

Adherent cells were cultured on 10 or 15 cm plates in a humidified incubator kept at 37 °C and 5% CO_2_. HEK293T cells were obtained from ATCC and cultured in Dulbecco’s Modified Eagle’s Medium (DMEM) supplemented with 10% fetal bovine serum (FBS) and 1% penicillin/streptomycin. Maintenance and subculturing of cells was conducted in accordance with ATCC guidelines.

### Plasmid Generation

All plasmids were generated by Gibson assembly cloning or golden gate cloning. For Gibson assembly, relevant DNA segments were amplified from plasmid templates by PCR with Q5^®^ High-Fidelity 2X Master Mix (NEB), and PCR products were separated by gel electrophoresis on a 1% agarose gel run at 120 V for 30 min then gel extraction with the Zymoclean Gel DNA Recovery Kit (Zymo Research) according to manufacturer’s protocol. PCR products were pooled in amounts according to manufacturer’s protocol and assembled with NEBuilder^®^ HiFi DNA Assembly 2x Master Mix (NEB).

For restriction cloning, the relevant ALTER backbone was linearized with BbsI-HF (NEB) and purified by gel electrophoresis and extraction, as described above. Primers were designed were annealed at a concentration of 10 µM each. The annealing reaction was diluted 1:10 and then ligated with T4 Ligase (NEB) according to the manufacturer’s protocol.

1 µl of the Gibson assembly or ligation mixture was added to a 10 µl aliquot of Stellar chemically competent cells (Takara) and incubated on ice for 1 h. Cells were transformed according to manufacturer’s protocol then plated onto an agarose plate with 100 µg/ml ampicillin. The plate was then incubated at 37 °C overnight to allow colony formation.

5 ml cultures of LB broth containing 100 µg/ml ampicillin were inoculated with single colonies and incubated at 37 °C overnight. Bacteria were pelleted by centrifuging at 3000×g for 5 minutes. Plasmids were purified from bacteria with the Quick-DNA Miniprep Plus Kit (Zymo Research).

Representative plasmid sequences are included in the Supplementary Information.

### Stable Cell line Generation with Sleeping Beauty Transposase

Gene cassettes for stable integration were cloned into plasmids flanked by Sleeping Beauty transposase recognition sites.^75^ Representative sequences are included in the Supplementary Information. On day 0, a 6-well cell culture plate was seeded with 300,000 HEK293T cells per well. On day 1, cells were transfected with jetPRIME transfection reagent in a transfection mixture containing: 2 µg plasmid, 0.2 µg Sleeping Beauty transposase plasmid (Addgene 34879), 4 µl jetPRIME reagent, 200 µl jetPRIME buffer according to manufacturer’s protocol. On day 2, media was exchanged. On day 3, the cells were lifted with trypsin and re-plated into 10 cm plates with 10 ml total media volume containing 250 µg/ml hygromycin or 2 µg/ml puromycin. When the cells reached 80–100% confluency, they were subcultured at a 1:20 ratio into a new 10 cm plate with 10 ml of antibiotic containing media. Untransfected controls were used to confirm selection efficacy. In total, cells were subject to selection for approximately 10 days.

For Tet-On-inducible cell lines, after selection, cells were plated into a 6-well culture dish at a density of 300,000 cells/well with 1 µM doxycycline. After two days of treatment with 1 µM doxycycline, cells were monoclonally sorted via FACS into a 96-well culture plate, selecting for cells in the top 10% of iRFP expression. Sorted cells which formed colonies were subcultured 1:2 into two wells of a 24-well plate and one well was treated with 10 µM doxycycline. Two days later, cells from the treated well were lifted by pipetting with PBS and analyzed by flow cytometry. Monoclonal lines which showed uniform induction of iRFP expression were selected and cells from the untreated wells were cultured as normal.

### Assessment of EGFP^Y66H^ Editing by Flow Cytometry

On Day 0 HEK293T cells stably expressing EGFP^Y66H^ were plated into a 24-well plate at a density of 70,000 cells/well. On Day 1cells were transfected with plasmids encoding an RNA editor and guide RNA using jetPRIME in accordance with the manufacturer’s protocol. Two days later, cells were lifted by pipetting with PBS and analyzed by flow cytometry.

For siRNA guides applied in HEK293T cells stably expressing EGFP^Y66H^ and an inducible editor, EGFP^Y66H^ experiments were carried out as above but 10 µM doxycycline was included in both the seeding and transfection media. 100 nM of siRNA was transfected using jetPRIME in accordance with the manufacturer’s protocol.

For ddEGFP^45^ knockdown experiments, HEK293T cells stably expressing ddEGFP were used in the same procedure.

### Assessment of Target Transcript Editing Rates and Expression Levels

HEK293T cells were plated into 6-well plates at a density of 250,000 cells/well. One day later, cells were transfected with a plasmid encoding an RNA editor and guide using jetPRIME transfection reagent in a transfection mixture containing: 3 µg plasmid, 6 µl jetPRIME reagent, 200 µl jetPRIME buffer. Transfection was carried out according to the manufacturer’s protocol. For experiments measuring editing on EGFP^Y66H^ transcripts cells were instead transfected with a 1:20 mixture of plasmids encoding the EGFP^Y66H^ driven by a PGK promoter and the editor/guide plasmid. Two days later, media was aspirated off the cells which were then washed with PBS and lifted by trypsin at room temperature. When cells had lifted, the trypsin was quenched with complete DMEM media, and the cell suspension was transferred to a 1.5 ml tube. Cells were pelleted by centrifugation at 300×g for 5 min and the supernatant media was aspirated. Cells were resuspended in 1 ml PBS then pelleted and the supernatant was aspirated.

Total RNA was extracted with either the RNEasy Kit (Qiagen) or Trizol (Thermo Fisher) according to the manufacturer’s protocol. RNA samples were eluted or resuspended with 40 µl DI water. RNA concentration was assessed by diluting samples 1:5 in pH=7.0 10 mM TrisCl buffer and taking the average of triplicate concentration measurements by nanodrop. RNA integrity was assessed by 1:5 dilution in pH=7.5 10 mM TrisCl buffer and confirming A260/280 2.0–2.1 and A260/230 > 2.0 by nanodrop. Furthermore 1.5 µl of each sample was separated on a 1% agarose gel, run at 60 V for 40 min, and assessed for clear 28S and 18S bands; when RNA extraction was performed with trizol the 5S subunit was also clearly visible.

For reverse transcription to cDNA, a 5 µl aliquot of each sample was diluted to a concentration of 125 ng/µl. 20 µl reverse transcription reaction mixtures included: (1) 4 µl 125 ng/µl total RNA (500 ng total); (2) 2 µl 10X RT buffer; (3) 1.4 µl 25 mM MgCl_2_; (4) 1 µl 10 mM dNTP mix; (5) 1 µl 100 mM DTT; (6) 1µl recombinant RNase inhibitor; (7) 1 µl, 50 µM oligoDT(16) primers (IDT); (8) 1 µl MultiScribe Reverse Transcriptase; (9) 7.6 µl DI water. The reverse transcription reaction was carried with the following protocol: (1) 25 °C, 10 min; (2) 37 °C, 2 h; (3) 95 °C; 5 min. When the reaction was complete cDNA samples were adjusted to a total volume of 100 µl by adding 80 µl of DI water.

All individual qPCR reactions were 20 µl total volume consisting of: (1) 10 µl SYBR™ Green Universal Master Mix (ThermoFisher); (2) 5 µl cDNA template; (3) 1.2 µl, 5 µM forward primer; (4) 1.2 µl, 5 µM reverse primer; (5) 2.6 µl DI water. For each experiment a master mix for each primer pair containing PCR master mix, primers, and water was made then aliquoted to individual reactions. Reactions were mixed in 96-well PCR plates sealed with Microseal B adhesive seals (BioRad) then analyzed by qPCR with the following protocol: (1) 50 °C, 2 min; (2) 95 °C, 2 min; (3) 95 °C, 15 s; (4) 60.1 °C (primer T_m_), 15 s; (5) 72 °C, 60 s/kilobase longest product; (6) fluorescence measurement; (7) repeat 3–6 39 times.

All qPCR primers were designed to amplify the edited region of the target transcript, with Primer Blast (NCBI), and appended with indexing adaptors. Primers were assessed for PCR efficiency using an 8-point dilution curve of pooled cDNA samples, run in technical duplicate or triplicate. Primer pairs were judged efficient if they resulted in apparent efficiencies of 90–110% over at least 5 points of the dilution curve. Primers were assessed for single product formation by melting curve (70–95 °C, +0.2 °C/5 s) and agarose gel electrophoresis. All primers used for qPCR in this study were efficient at a T_m_ of 60.1 °C.

Final qPCR measurements were carried out in technical triplicate for the target transcript and two reference transcripts, SNW1 and GAPDH or NHNRPL.^76^ Final qPCR data was analyzed according to the method recommended by Taylor et al.^77^

For editing assessment by sequencing, a PCR amplification number, *n*, was identified such that all samples were at 50–75% maximum amplification at *n* cycles, based on qPCR data. Edited segments of target transcripts were amplified by PCR using Amplitaq Gold 360 2X Master Mix A 10 µl aliquot of the PCR reaction was assessed by agarose gel electrophoresis and the remainder was purified with Zymo DNA Clean & Concentrator kit (Zymo Research) according to the manufacturer’s protocol. An aliquot of the purified PCR product was diluted 1:100 and amplified by 12-cycle PCR with primers containing barcode sequences. PCR products were assessed and purified as above, then pooled in equal amounts in groups of three (usually 3 replicates of the same condition) and sequenced by Oxford nanopore sequencing. Reads in the fastq sequencing data were sorted by barcode. Adaptors and low quality bases were trimmed with cutadapt,^78^ and editing was quantified with CRISPResso2.^79^ See Supplementary Information for code and software argument details.

### RNAseq Total RNA Extraction

HEK293T cells were plated into 6-well plates at a density of 250,000 cells/well. One day later, cells were transfected with a plasmid encoding an RNA editor and guide using jetPRIME transfection reagent in a transfection mixture containing: 3 µg plasmid, 6 µl jetPRIME reagent, 200 µl jetPRIME buffer. Transfection was carried out according to the manufacturer’s protocol. Two days later, media was aspirated off the cells which were then washed with PBS and lifted by treatment with trypsin at room temperature. When cells had lifted, the trypsin was quenched with complete DMEM media, and the cell suspension was transferred to a 1.5 ml tube. Cells were pelleted by centrifugation at 300×g for 5 min and the supernatant media was aspirated. Cells were resuspended in 1 ml PBS then pelleted and the supernatant was aspirated.

Total RNA was extracted with Trizol (ThermoFisher) according to the manufacturer’s protocol. RNA samples were resuspended in 40 µl DI water. To remove residual DNA, the aqueous samples were treated with DNAse from the Qiagen RNAse-Free DNase Set and re-purified with the RNEasy Kit (Qiagen) according to the manufacturer’s instructions, final elution in 40 µl DI water. RNA concentration and integrity was assessed as above. Samples were stored at −80 °C then transferred to Novogene (see below) on dry ice by courier at the earliest opportunity. All data was collected from three biological replicates.

### RNAseq Sample Quality Control and Library Preparation

RNAseq data was obtained from Novogene via their mRNAseq service. According to their methods documents:

The concentration, purity, and integrity of total RNA samples was assessed with an Agilent 5400 bioanalyzer. Results for sample quality control can be found in the Supplementary Information.

Messenger RNA was purified from total RNA using poly-T oligo-attached magnetic beads. After fragmentation, the first strand cDNA was synthesized using random hexamer primers followed by the second strand cDNA synthesis. The library was ready after end repair, A-tailing, adapter ligation, size selection, amplification, and purification. The library was checked with Qubit and real-time PCR for quantification and bioanalyzer for size distribution detection. After library quality control, different libraries were pooled based on the effective concentration and targeted data amount, then subjected to 2×150 bp paired-end sequencing on an Illumina NovaSeq X Plus resulting in at least 21,223,588 paired end reads for sequenced samples.

### RNAseq Data Analysis General Information

Detailed argument information for all software tools and code for all Python and R scripts used in RNAseq data analysis can be found in the Supplementary Information. All data analysis was performed on personal computers or Stanford’s Sherlock computing cluster.

### RNAseq Data Quality Control

Fastq files with raw read data were obtained from Novogene for ALTER and from BioProject PRJNA635732^12^ for Cas editors and A3A overexpression. Trimming of sequencing adaptors and low quality bases, and filtering of low quality reads was performed with fastp.^80^ The resulting fastq files were assessed with fastqc and multiqc.^81^

### RNAseq Genomic References

All genomic reference data was derived from the GENCODE release 48 comprehensive gene annotation of the primary GRCh38.p14 assembly. For each sample the relevant editor plasmids were added to the genome fasta, with corresponding transcript sequences added to the transcript fasta and features added to the GENCODE gtf file. The genome fasta and gtf files were used to generate genome indexes for STAR.^82^

### Analysis of C-to-U Off Targets in RNAseq Data

Reads were aligned to the corresponding genomic reference using STAR^82^ v2.7.10b in two-pass mode. Variant calling was performed with GATK^83^ v4.6.0.0. Alignment bam files were prepared for variant calling. Duplicate reads were identified and annotated with GATK MarkDuplicates and reads spanning splice junctions were split with GATK SplitNCigarReads. Variants were called with GATK HapplotypeCaller in GVCF mode, parallelized over four genomic intervals per sample. GVCF files for each sample were merged with GATK MergeVcfs and the sample GVCF files were then combined with GATK CombineGVCF. The combined GVCF underwent joint genotyping with GATK GenotypeGVCFs to produce a final variant vcf file. This vcf file was filtered for low QD (QD<2.0), strand bias (FS>30.0), low depth (DP<10), and low quality (QUAL<20.0) then converted to a variant table with GATK VariantFiltration and GATK VariantsToTable. The data from Novogene and BioProject PRJNA635732^12^ were analyzed in separate groups.

Final analysis was carried out with Python scripts developed for this project. The variant table was processed by: (1) inferring variant strand and sequence identity, (2) cross referencing variants to the ClinVar^84^ database, (3) converting to binary genotypes, (3) calculating the percentage of reads matching the reference nucleotide, (4) calculating the percentage of reads matching the variant base for C-to-U variants. The final table was then filtered for high-confidence C-to-U off targets based on the following criteria: (1) called as variants in the editor sample, (2) C-to-U variant, (3) read depth >= 20, (4) genotype quality >= 20 (99% confidence), (5) >99% of transfection control reads match the reference base. Sequence contexts for off-target hits were extracted from genomic reference data and used to generate nucleotide frequency plots. The mfe secondary structure of each sequence context was calculated and used to generate structural element frequency plots. The ALTER guide sequence was aligned to each sequence context and alignment data was used to generate alignment frequency plots.

### Analysis of Differentially Expressed Genes in RNAseq Data

Using Salmon,^85^ raw transcript counts were generated by mapping reads from sequencing fastq files to a Salmon index generated from the genomic reference transcripts fasta file. Count normalization and DEG analysis were performed using DESeq2^86^ in an R script written for this project. Genes were identified as DEGs if padj <0.01 and the magnitude of log_2_(fold-change) > 1. A list of all genes with sufficient read depth to be included in the DESeq2 ranked by log_2_(fold-change) underwent GSEA analysis^60^ using the fgsea^87^ library in R. DEG hits were analyzed for GO term enrichment with the clusterProfiler^88^ R library. DEGs were cross-referenced to known miRNA targets in miRtarBase^61^ using a Python script.

### Editor Analysis by MARIA

Peptides were predicted for MHC presentation via a local copy of the publicly available tool MARIA.^62^ In brief, all possible 15mer peptides containing a non-native residue were predicted for a given sequence. These peptides are assigned a MARIA raw score and a percentile score corresponding to their predicted raw score relative to 20,000 random human peptides. Using previously determined percentile thresholds, peptides were considered predicted to be presented if in the 63rd percentile of MARIA raw scores, representing a false omission rate of 1%. Unless otherwise stated, predictions were performed using the HLA-DRB1*01:01 allele without the optional TPM parameter.

### Predicted Peptide Immunogenic Silencing

Protein sequences were analyzed to identify regions containing non-native residues, including engineered amino-acid substitutions, heterologous sequence insertions, and domain-junction boundaries. The full amino-acid sequence was scanned using overlapping 15mer tiles, cataloging all peptides containing at least one non-native residue. Overlapping peptides were merged into contiguous intervals, defining discrete immunogenic regions for targeted mutation.

For each region, single- or double-amino-acid substitutions were introduced at eligible positions to reduce predicted immunogenicity. Each extended sequence window was then evaluated for presented peptides as described above. Variant regions exhibiting the lowest predicted presented peptides were chosen with the total number of tied variants limited by random subsampling. These winning variants were then further mutated as described above until resulting in variant regions with zero predicted presented peptides. Resulting designs were advanced for subsequent experimental benchmarking. This analysis was carried out in an R script developed for this project.

### VLP Production

Information on plasmid sequences can be found in the Supplementary Information.

Day 0, a T75 culture flask was seeded with 3*10^6^ HEK293T cells in complete DMEM. On day 1, the cell culture media was exchanged for 10 ml of complete DMEM and the cells were transfected with the following plasmid mixture: (1) 6.7 µg pRFL396, encoding the Gag-ALTER-4 fusion; (2) 3.3 µg MMLVgag–pro–pol (Addgene #35614); (3) 1 µg VSV-G (Addgene #8454); (4) 10 µg pRFL364 encoding the ALTER guide to be expressed transiently in the producing cells. The plasmids were mixed with 500 µl jetPRIME buffer and 20 µl jetPRIME reagent (Sartorius) and transfection was carried out according to the manufacturer’s protocol. 6 h post-transfection the media was exchanged for 10 ml fresh complete DMEM further supplemented with 1X ViralBoost (Alstem). On day 2, the cell culture media was collected in a 15 ml tube and replaced with 10 ml fresh complete DMEM. The 15 ml tube was centrifuged at 500×g, 4 °C, 5 min, then the supernatant was transferred to a second tube. 2.5 ml PEG-IT (System Biosciences LV810A-1) was added to the sample and it was stored at 4 °C. On day 3, the cell culture media was collected in a 15 ml tube centrifuged at 500×g, 4 °C, 5 min, then the supernatant was transferred to a second tube. 2.5 ml PEG-IT was added to the sample and it was stored at 4 °C. On day 4, the samples were centrifuged at 1500×g, 4 °C, 30 min. The supernatant was transferred to a new tube, carefully as not to disturb the precipitant pellet. The supernatant was again centrifuged at 1500×g, 4 °C, 30 min and the supernatant was aspirated away from the residual pellet. All pellets (24 h primary, 24 h residual, 48 h primary, 48 h residual) were mixed by resuspending in 200 µl PBS then stored at −80 °C in 50 µl aliquots.

### Assessment of VLP Activity by EGFP^Y66H^ Editing

On day 0, a 24-well culture plate was seeded with 25,000 HEK293T cells stably expressing the EGFP^Y66H^ reporter. On day 1, the complete DMEM media was replaced with 0.5 ml fresh complete DMEM. One well was treated with a 50 µl aliquot of the VLP suspension while another was transfected with 500 ng plasmid encoding ALTER-4 and guide with jetPRIME reagent (Sartorius) according to the manufacturer’s protocol. On day 5, the cells were analyzed by flow cytometry.

### AAV Production

On day 0, HEK293T cells were plated into 9 6-well plates per viral sample at a density of 350,000 cells/well. Day 1, cells were transfected with transgene, AAV-DJ rep/cap (Addgene #130878), and helper plasmid (GenBank AF369965.1) in a 1:1:2 ratio using jetPRIME transfection reagent in a transfection mixture containing: 2 µg plasmids, 4 µl jetPRIME reagent, 200 µl jetPRIME buffer. Transfection was carried out according to the manufacturer’s protocol. Day 4, media was aspirated off the cells which were then washed with 350 µl PBS and lifted by treatment with 200 µl trypsin at room temperature. When cells had lifted, the trypsin was quenched with 800 µl complete DMEM media. The cell suspension for all 9 wells was pooled into a 15 ml conical tube and pelleted by centrifugation at 500×g for 5 min. The supernatant media was aspirated, and the pellet was resuspended in 750 µl complete DMEM media. Cell suspensions where then transferred to cryovials and lysed doing three freeze thaw cycles going from dry ice in ethanol to a 37 °C water bath, 3 min in each per cycle. Lysates were centrifuged at 7,500×g for 5 min. The supernatant was extracted and centrifuged like before. If sediment was detected after extracting the supernatant, the supernatant was spun a final time at 9,000×g for 5 min. Supernatant was used for immediate transduction into HEK293T cells stably expressing EGFP^Y66H^.

### AAV Titer Measurement

200 µl of viral containing supernatant was treated with benzonase nuclease (MilliporeSigma) at 50U/mL to remove uncapsidated viral genomes. The mixture was incubated at 37 °C for 45 min, after which the enzyme was inactivated by addition of UltraPure™ 0.5M EDTA (ThermoFisher) to a final concentration of 15 mM. The Macherey-Nagel NucleoSpin virus mini kit (Fisher Scientific) was used to isolate viral DNA from the mixture and used according to the manufacturer’s protocol. Transgene plasmid was linearized, and concentration was assessed by nanodrop. Six qPCR standards were prepared by serial 10-fold dilutions starting from a 5 ng/µL stock of the linearized plasmid. Viral DNA was diluted 10-fold and measured against the standards by qPCR with primers against the transgene.

### Assessment of AAV Activity by EGFP^Y66H^ Editing

A day before AAV harvesting, day 0, HEK293T cells stably expressing EGFP^Y66H^ under an EF1α promoter were plated into 48-well plates at a density of 42,500 cells/well. Day 1, media was aspirated off the cells and replaced with a mixture of 190 µl complete DMEM and 60 µl AAV suspension. Day 5, media was aspirated off and the cells washed with PBS. Cells were then lifted by treatment with trypsin at room temperature. When cells had lifted, the trypsin was quenched with complete DMEM media, and cells were pelleted by centrifugation at 500×g for 5 min then resuspended in PBS. After pelleting by centrifugation at 500×g for 5 min, cells were resuspended in PBS and analyzed by flow cytometry.

